# Effect of hydrogen inhalation on IL-40 and SIgA in a Rat Model of Pulmonary Mucosal Immunity

**DOI:** 10.1101/2020.06.29.177345

**Authors:** Yiping Ma, Zhu Li, Yalei Zhao, Mo Sun, Wuzhuang Sun, Jiechao Wang

**Affiliations:** Department of Respiratory Medicine, The First Hospital of Hebei Medical University, Shijiazhuang, China; Department of General Surgery, The Fourth Hospital of Hebei Medical University, Shijiazhuang, China; Department of Radiotherapy, The Fourth Hospital of Hebei Medical University, Shijiazhuang, China; Deparment of Vasculocardiology, Hebei Geriatric Hospital, Shijiazhuang, China

**Keywords:** hydrogen, SIgA, IL-40, TGF-β1, COPD

## Abstract

**Background:** Recently, some researchers have reported that PIgR expression is down-regulated in Chronic Obstructive Pulmonary Disease (COPD) and SIgA deficiency correlates with severity of airflow obstruction. What’ s more, some studies have demonstrated that 2 percent of hydrogen or hydrogen water is effective in treating and preventing various diseases.

**Objectives:** The aim of this study was to observe the effect of hydrogen on the expression of SIgA, PIgR, IL-4, IL-5, TGF-β1 and IL-40 in lung tissue of COPD rats, to study the relationship between lung pathology parameter and SIgA, PIgR, therefore we can understand the effect of hydrogen on the development of COPD by changing SIgA expression of airway mucosal in COPD rats.

**Methods:** A rat model of COPD was established by cigarette smoke exposure, and different concentrations of hydrogen were inhaled as intervention measures. After 4 months of cigarette smoke exposure, pathologic changes and airway wall remodeling of the lung were assessed by optical microscope. The protein expressions of SIgA, PIgR, IL-4, IL-5, TGF-β1 as well as IL-40 in the lung tissues were observed by immunohistochemistry or Western blot. The correlation between lung pathology parameter and the expression of SIgA, PIgR was analyzed. The correlation between SIgA and the expression of IL-4, IL-5, TGF-β1 and IL-40 was analyzed.

**Results:** The results showed that hydrogen inhalation significantly ameliorated lung pathology and airway wall remodeling, increased the protein expression of SIgA, PIgR, IL-4, IL-5, and IL-40, and reduced the protein expression of TGF-β1.

**Conclusions:** Inhalation of 22% and 41.6% hydrogen showed a better effect than inhalation of 2% hydrogen. Hydrogen inhalation can significantly improve the expression of SIgA on the mucosal surface of COPD rats, which may be one of the mechanisms which hydrogen works on COPD pathogenesis.

## Introduction

Chronic Obstructive Pulmonary Disease (COPD) is a common, preventable and treatable disease, which is characterized by irreversible airflow limitation due to airway and/or alveolar abnormalities, usually caused by significant exposure to harmful particles or gases(1, 2). It is currently reported to be the fourth leading cause of death worldwide, and also expected to be the third leading cause of death by 2020(3). Recently, some researchers have found that mucosal deficiency is related to chronic airway inflammation. Subsequently, the study has shown that local IgA deficiency is associated with bacterial translocation, small airway inflammation, and airway remodeling(4).

Surface IgA (SIgA), along with mucociliary clearance, defenses against inhaled antigens and microorganisms by preventing airway inflammation(5). In small airways, IgA is produced locally by subepithelial plasma cells. A dimer of two IgA monomers linked by J(“joining”) chain and secretory component (SC) composes SIgA(6), which is transported across airway epithelial cells via the polymeric immunoglobulin receptor (pIgR). After transportion to the mucosal surface, pIgR is cleaved and SC remains attached to form SIgA(7). SC is an extracellular component of pIgR, which plays a role in stabilizing the structure of SIgA in the process of transportion(8). Therefore, SIgA and SC play a pivotal role in the immunity of respiratory tract(9, 10). In 2001, Pilette et al reported that SC expression was reduced in airways of COPD(11). Subsequently, researchers found that patients with COPD had reduced SIgA in bronchoalveolar lavage(12). Polosukhin et al have shown that pIgR expression is down-regulated in COPD and that SIgA deficiency is correlated with severity of airflow obstruction(13). Richmond et al have documented that, as a consequence, pIgR deficiency leads to chronic neutrophil inflammation, which contributes to small airway fibrosis and emphysema(14). Together with what has been published above, dysfunction of bronchial epithelial mucosa led to abnormalities in SIgA transported to the airway, which impairs the defense ability of local hosts. Therefore, persistent structural and functional abnormalities of airway epithelium play a key role in development and progression of COPD.

Cytokine, acts as a secreted protein, may plays a vital role in host defense, inflammation, and immune responses. In 2017, Catalan et al described the latest cytokine, IL-40, which was found to be expressed by peripheral B cells upon activation, and encoded a small secreted protein(27kDa) of 265 amino acids by an uncharacterized gene(15). What’ s more, they also found IL-40-/-mice exhibited smaller Peyer’ s patches, lower numbers of IgA-secreting cells, and fewer IgA in the serum, gut, feces, and lactating mammary gland(16). Interestingly, TGF-β1 had been shown to play an important role in potentiating the expression of IL-40 by activated B cells, and a progressive increase was observed when B cells were treated with IL-4 and TGF-β1. There was a direct correlation between the expression of IL-40 and the onset of IgA production. Understandably, IL-40 was involved in the development of humoral immune responses, especially those associated with IgA production.

Together, our researches shed light on the potential mechanism of SIgA expression in COPD rats. In recent years, a number of studies have shown that hydrogen therapy has a positive effect on various diseases(17–20). Liu et al in our team found that different concentrations of hydrogen inhalation could significantly reduce inflammation and apoptosis, protect mitochondria, and ameliorate cardiovascular function in COPD rat model(21). A key question is whether there exist a possibility that hydrogen inhalation will have a positive effect on SIgA expression in rat model of COPD, which is to be tested in the experiment.

## Materials and methods

### Instruments and reagents

A smoking device manufactured by Shijiazhuang Jinyang Science and Technology Development Co., Ltd. (JY-01 model, Shijiazhuang, Hebei, China) and Cigarette China Diamond purchased from Hebei Tobacco Company (tar: 13 mg, nicotine: 1.2 mg, flue gas carbon monoxide: 14 mg, Zhangjiakou, Hebei, China) were used for producing the COPD model. A hydroxide atomizer manufactured by Shanghai Asclepius Meditec Co., Ltd. (AMS-H-01, Shijiazhuang, Hebei, China) was used in this study. Nitrogen (15% O_2_, 85% N_2_) was purchased from Shijiazhuang Central Plains Specialty Gases Ltd., and a 2% H_2_ gas mixture (21% O_2_, 2% H_2_, 77% N_2_) was purchased from Guangzhou Puyuan Gas Company Ltd. Anti-IgA secretory component (Mouse, ab212330, 1:500 in IHC, Abcam), anti-IL-4 (Mouse, sc53084, 1:100 in IHC, 1:500 in WB; SanTaCruz), anti-PIGR (Rabbit, bs-6061R, 1:500 in WB; Bioss), anti-IL-5 (Rabbit, GTX55678, 1:100 in IHC, 1:500 in WB; GeneTex), anti-TGF-β1 (Rabbit, ARG56429, 1:500 in WB; Arigo), anti-C17orf99 (Rabbit, PA5-71295, 1:500 in WB; ThermoFisher) primary antibodies, and anti-rabbit and anti-mouse secondary antibodies (Abbkine, Reddlands, CA, USA) were used to determine protein expression.

### Animal model

Healthy male Sprague-Dawley rats aged 7 weeks (body weight 180±20g), were obtained from Animal Experimental Center of Hebei Medical University. Prior to the experiment, animals were accommodated for 1 week. The rats were housed in a centralized animal room and provided with a standard diet and water ad libitum. All animal handling protocols and experimental procedures conformed to Regulations of Beijing Municipality on the Administration of Laboratory Animals. This study was approved by the Institutional Animal Care and Use Committee of Hebei Medical University.

An animal model was established by the smoking method. All animals were randomly divided into 5 groups (10 rats in each group): control group, COPD group, high-hydrogen group (Hh group, which was exposed to 21% O_2_, 41.6% H_2_, and 100% N_2_ at a flow rate of 5:3), intermediate hydrogen group (Hm group, which was exposed to 21% O_2_, 22% H_2_, and 15% O_2_, 85% N_2_ at a flow rate of 1:2), and low-hydrogen group(Hl group, which was exposed to 21% O_2_, 2% H_2_). In addition to the control group, cigarette smoke and hydrogen were in reality exposed for 4 months. In the meantime, the control group received no intervention during 4-month period.

### Tissue Preparation

Tissue preparation was performed as described previously(22). The lung was harvested and fixed immediately in buffered 4% paraformaldehyde for 24 h. The tissue was subsequently dehydrated in a graded ethanol series and embedded into paraffin. For histomorphology analysis, consecutive 5-μm lung sections were collected. Sections were prepared for Hematoxylin and eosin (HE) staining, Masson’s trichrome staining, and immunohistochemical staining and examined under an optical microscope (Olympus IX71; Olympus, Tokyo, Japan).

### HE staining

HE staining was performed by standard protocol. For routine histologic examination, the 5μm sections were stained. Pathologic changes in the lung tissues were observed under an optical microscope. MLI was determined for each region studied on an overlay consisting of horizontal and vertical lines. MAN was determined according to alveolar number in each field of view and a square area of the field.

### Masson’s trichrome staining

Masson’s trichrome staining was performed by standard protocol. The value of bronchial airway was selected which was defined as the difference between the area delineated by the inner boundary of tunica intima (A1) and the external boundary of tunica adventitia (A2), divided by the length of the external boundary of tunica intima. The area of the wall (A3) was calculated with A1 and A2 (A3 = A2 - A1), and the ratio of wall and lumina was achieved with A3 and A2 (the ratio = A3/A2). The bronchial wall thickness/ diameter (BWT/D) ratio was determined.

### Immunohistochemistry for the Detection of SIgA, IL-4 and IL-5 in Pulmonary Tissue

The protein expression levels of SIgA, IL-4 and IL-5 were assessed by IHC. Before using microwave antigen retrieval, deparaffinized sections were pretreated, followed by incubation in 3% H_2_O_2_ at 25°C for 25 min, and goat serum at 37°C for 40 min. Next, the tissues were incubated respectively with primary antibodies overnight at 4°C. After rinsed with PBS, the tissues wereincubated with biotinylated secondary antibody at 37°C for 40 min and subsequently with horseradish peroxidase (HRP)-conjugated biotin at 37°C for 40 min. Finally, diaminobenzidine (DAB) was used for chromogenic reaction, and then hematoxylin dyeing was performed for visualizing locations in the sections. The average number of positive indicators was calculated in view fields.

### Western Blot (WB) for Analysis of PIgR, IL-4, IL-5, TGF-β1 and IL-40 in Pulmonary Tissue

The protein expression levels of PIgR, IL-4, IL-5, TGF-β1 and IL-40 in the tissues were measured using WB. Firstly, lung tissues were collected and lysate was added. Quantify total protein concentration was determined by the Lowry method. The protein were separated by corresponding SDS-PAGE, and then electrophoresed and transferred to a nitrocellulosemembrane. After blocking in skim milk, the membranes were incubated with the primary antibody at 4°C for overnight. According to the primary antibody, the membranes were incubated with secondary antibody at 25°C for 1h. The chemiluminescence imaging analyzer was used for scanning imaging. Each experiment was repeated five times, and similar results were obtained from the chemiluminescence imaging analyzer.

### Statistical Analysis

Data were presented as the mean±standard deviation (SD). Student-Newman-Keuls method was used for Pairwise comparisons with homogeneity of variance and one-way analysis of variance (ANOVA) was applied for comparison among groups. Correlation analysis of two sets of normal distribution measurement data using Pearson linear correlation analysis, and further using linear regression analysis to fit the equation. *P*<0.05 was considered statistical significance. All experimental data were analyzed by SPSS software 21.0.

## Results

### The pathological changes of lung tissues in COPD rats with hydrogen inhalation

In control group, the alveolar structure was normal, the trachea mucosal was intact, pulmonary artery wall structure was normal, and no inflammatory cell infiltration was found. In COPD group, alveolar duct, alveolar vesicle and alveoli were significantly enlarged, alveolar structure was disordered, alveolar wall was broken, pulmonary bullae was partial fusion, ciliated cell were degeneration and necrosis, the airway wall thickness increased obviously, the airway mucosal was damaged seriously, and the airway wall remodeling increased obviously, pulmonary arteriole wall was thickening, and different levels of infiltration by neutrophils, lymphocytes and monocytes were observed.

Compared with COPD group, the thickening of alveolar wall, airwway wall and pulmonary artery wall was ameliorated, the condition of alveolar fracture was better, inflammatory cell infiltration and the number of bullae was less in Hh, Hm and Hl groups (Fig 1A).

**Fig 1.**
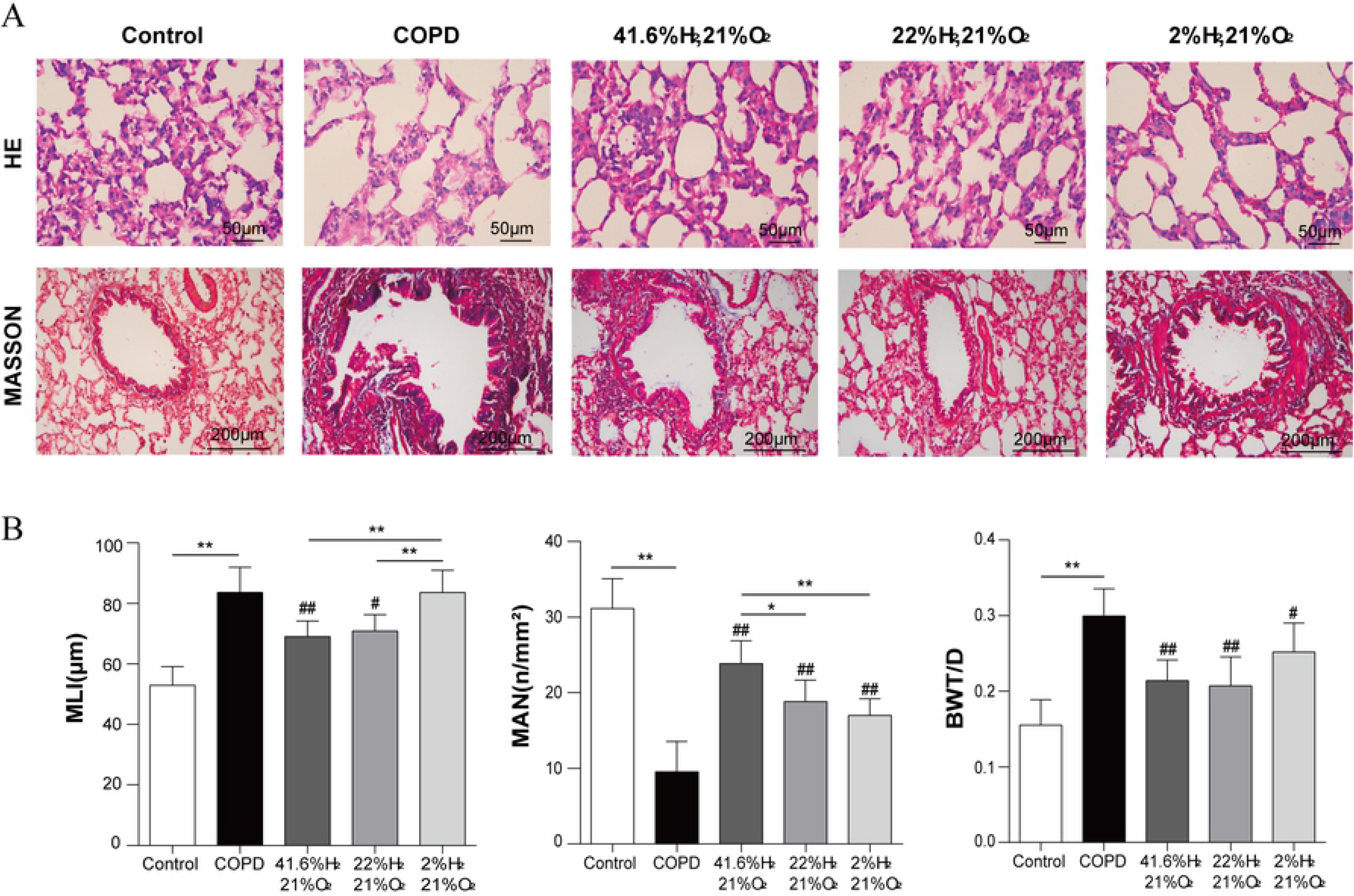
Effect of hydrogen on the lung pathologic changes and airway remodeling in the COPD rat model. **A** Rat lung pathologic observations (HE×400), Bronchial airway remodeling observations (MASSON×200). **B** MLI, MAN, and BWT/D were measured to compare the effect of hydrogen in the COPD rat model. (n=10 for each group. ^##^*P*<0.01,^#^*P*<0.05 compared with COPD group; \**P*<0.05,\*\**P*<0.01).

Compared with control group, MLI was significantly increased (*P*<0.01), MAN was significantly decreased in COPD group (*P*<0.01). Compared with COPD group, MLI was decreased in Hh group and Hm group (*P*<0.01, *P*<0.05 respectively). Compared with Hl group, MLI was significantly lower in Hh group and Hm group (*P*<0.01). Compared with COPD group, MAN was significantly increased in Hh, Hm, Hl groups (*P*<0.01). Compared with Hh group, MAN was decreased in Hm group and Hl group (*P*<0.01, *P*<0.05 respectively), but there was no significant difference between Hm group and Hl group (*P*>0.05). Compared with control group, BWT/D was significantly increased in COPD group (*P*<0.01). Compared with COPD group, BWT/D was reduced in Hh, Hm and Hl groups (*P*<0.01, *P*<0.05 respectively) (Fig 1B).

### The expression of SIgA, IL-4 and IL-5 in lung tissue was detected by immunohistochemistry

Compared with control group, the expression of IL-4, IL-5 was reduced in COPD group. In contrast with COPD group, IL-4, IL-5 was increased in Hh group and Hm group. Compared with COPD group, IL-4, IL-5 show no difference in Hl group (Fig 3A). Compared with control group, SIgA was significantly reduced in COPD group (*P*<0.01). Compared with COPD group, SIgA was significantly increased in Hh group and Hm group (*P*<0.01), there was no significant difference in Hl group (*P*>0.05), but there was no significant difference between Hh group and Hm group (*P*>0.05) (Fig 2A, 2B).

**Fig 2.**
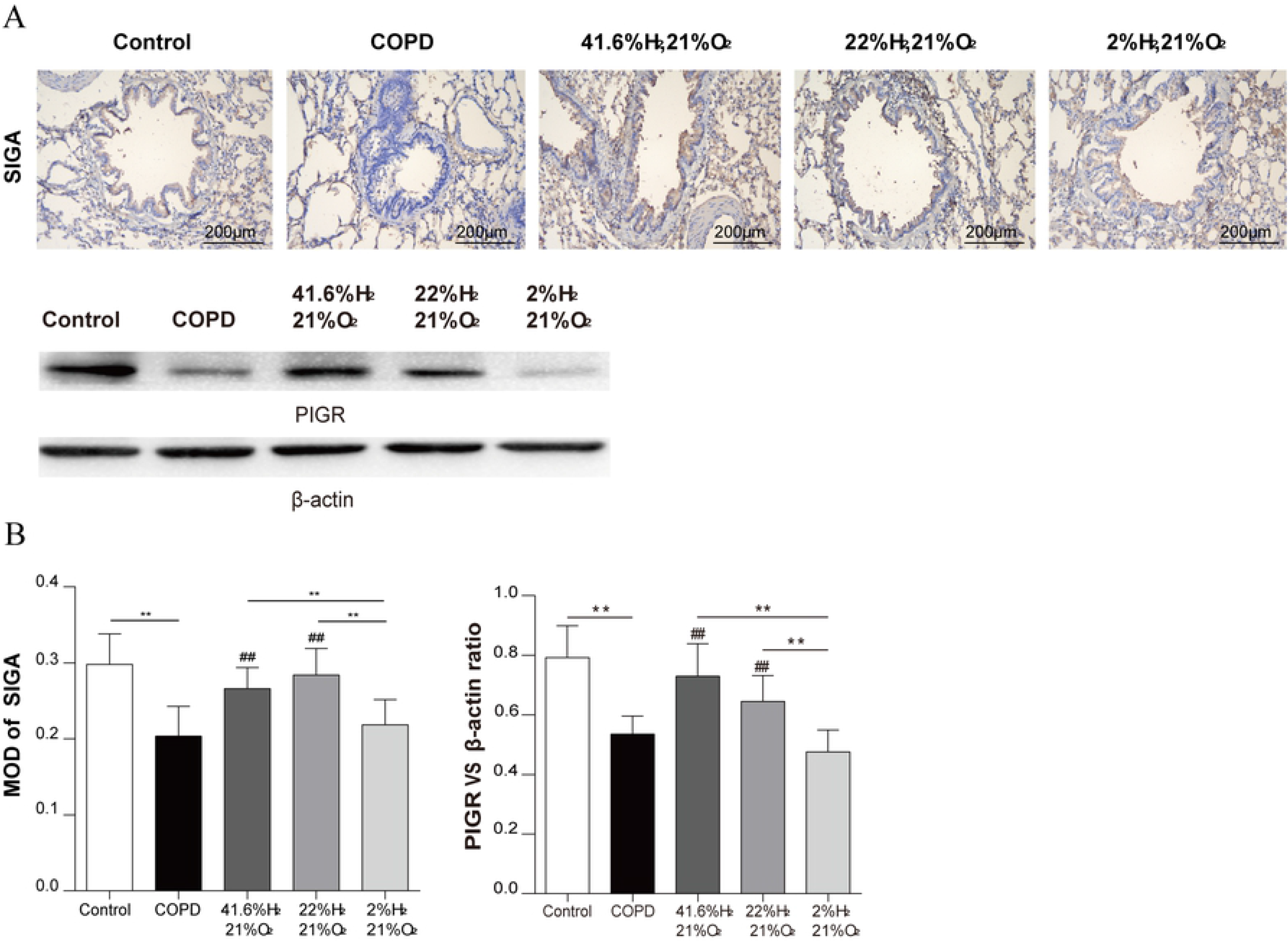
Effect of hydrogen on the expression of SIGA and PIgR in the lung obtained from the COPD rat model, determined by immunohistochemical staining or Western blotting. **A** Immunohistochemical staining of the lung sections for SIGA, and Western blotting of the lung sections for PIgR. **B** Quantitative analysis of the expression of SIGA and PIgR in the COPD rat model. (n=10 for each group. ^##^*P*<0.01 compared with COPD group; \*\**P*<0.01). Each experiment was repeated three times and similar results were obtained.

**Fig 3.**
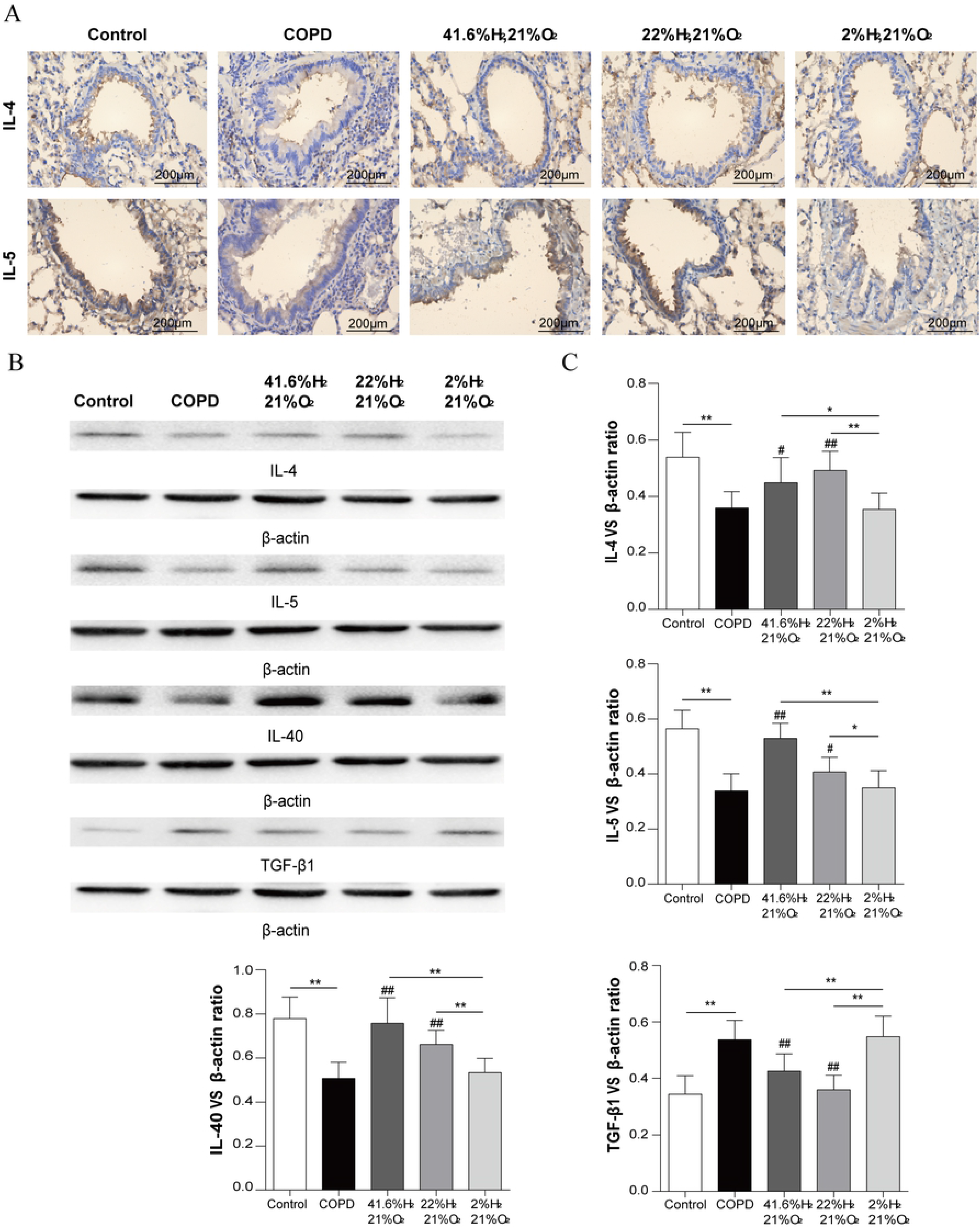
Effect of hydrogen on the expression of IL-4, IL-5, TGF-β1 and IL-40 in the lung obtained from the COPD rat model, determined by immunohistochemical staining or Western blotting. **A** Immunohistochemical staining or Western blotting of the lung sections for IL-4, IL-5, TGF-β1 and IL-40. **B** The target protein bands were densitometrically. **C** Quantitative analysis of the expression of IL-4, IL-5, TGF-β1 and IL-40 in the COPD rat model. (^#^*P*<0.05, ^##^*P*<0.01 compared with COPD group; \**P*<0.05,\*\**P*<0.01). Each experiment was repeated three times and similar results were obtained.

### The expression of PIgR, IL-4, IL-5, TGF-β1 and IL-40 in lung tissue was detected by Western blot

Compared with control group, the expression of IL-4, IL-5, IL-40, PIgR was significantly reduced (*P*<0.01), TGF-β1 was significantly increased in COPD group (*P*<0.01); Compared with COPD group, the expression of IL-4, IL-5, IL-40, PIgR in lung tissue was increased (*P*<0.01, *P*<0.05 respectively), TGF-β1 was significantly reduced in Hh group and Hm group (*P*<0.01); Compared with COPD group, the expression of IL-4, IL-5, TGF-β1, IL-40, PIgR shows no difference in Hl group (*P*>0.05) (Fig 2A, 2B, 3B, 3C).

### Correlation between SIgA, PIgR and BWT/D

SIgA was negatively correlated with BWT/D (r=−0.5839, *P*<0.01), PIgR was negatively correlated with BWT/D (r=−0.6051, *P*<0.01) (Fig 4A).

**Fig 4.**
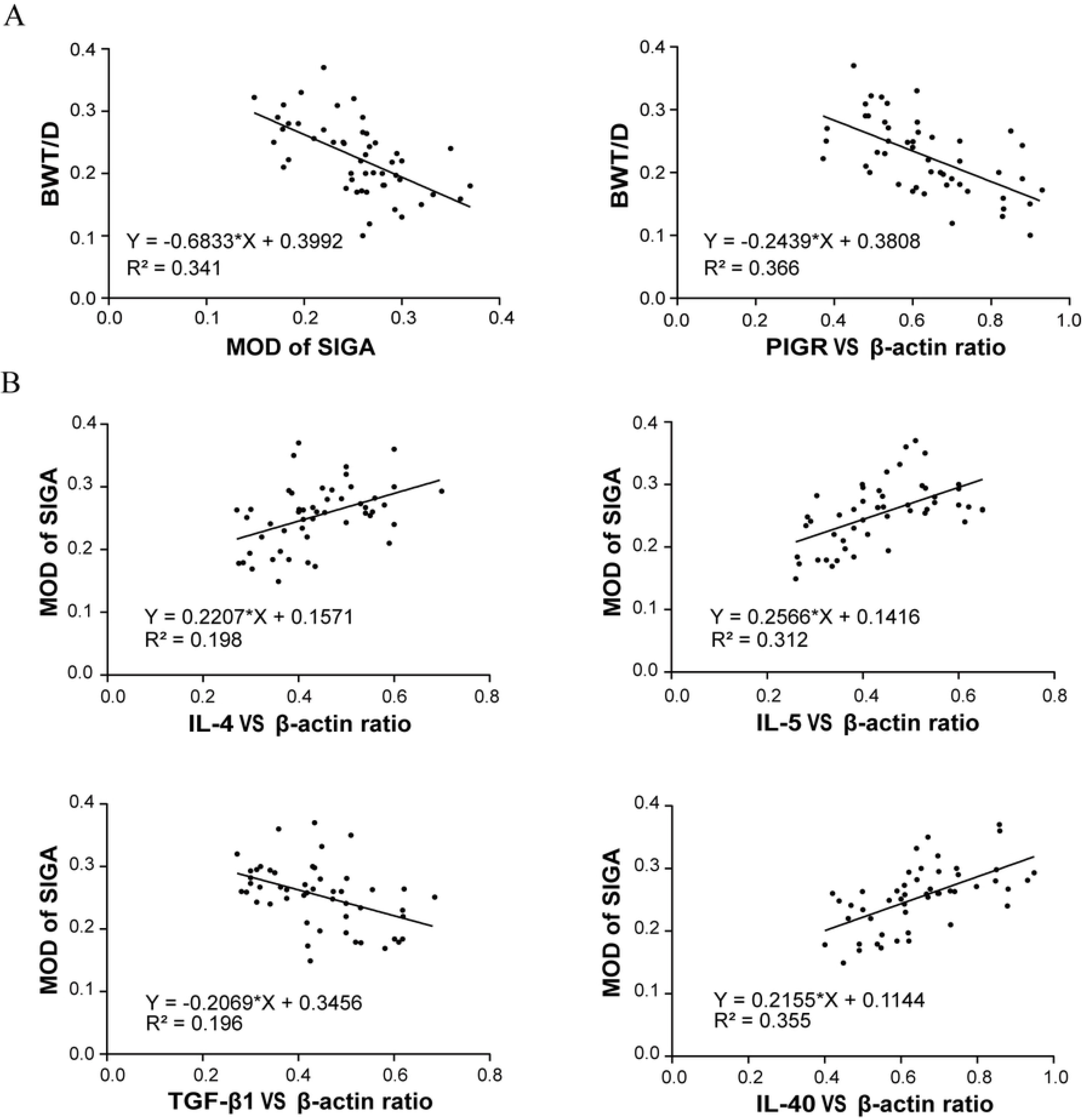
Correlation of comparative SIgA、PIgR and BWT/D; Correlation of comparative IL-4, IL-5, TGF-β1, IL-40 and SIgA. **A** Correlation of comparative MOD of SIGA to BWT/D; Correlation of comparative PIGR vs β-actin ratio to BWT/D. **B** Correlation of comparative IL-4 vs β-actin ratio to MOD of SIGA; Correlation of comparative IL-5 vs β-actin ratio to MOD of SIGA; Correlation of comparative TGF-β1 vs β-actin ratio to MOD of SIGA; Correlation of comparative IL-40 vs β-actin ratio to MOD of SIGA.

### Correlation between SIgA and IL-4, IL-5, TGF-β1, IL-40

IL-4 vs β-actin ratio was positively correlated with MOD of SIgA (r=0.4449, *P*<0.01); IL-5 vs β-actin ratio was positively correlated with MOD of SIgA (r=0.5589, *P*<0.01); TGF-β1 vs β-actin ratio was negatively correlated with MOD of SIgA (r=−0.4422, *P*<0.01); IL-40 vs β-actin ratio was positively correlated with MOD of SIgA (r=0.5959, *P*<0.01) (Fig 4B).

## Discussion

Bronchial epithelial SIgA is the critical antibody for respiratory immune defense against inhaled antigens and microorganisms at mucosal surfaces, including the airways. Recently, studies have shown that SIgA deficiency is widespread in COPD. Therefore, how to improve the bronchial mucosal immunity is a core issue. Our previous study showed that hydrogen inhalation could visibly reduce inflammation and apoptosis, and improve cardiovascular function, suggesting a “significant effect” of hydrogen inhalation on COPD(21). Our results provided new information that treatment with hydrogen could ameliorate pathological changes, airway remodeling as well as SIgA expression. Moreover, we found that Hm hydrogen concentration was necessary for SIgA expression. Together, our findings pinpointed a positive role of hydrogen inhalation in improving mucosal immunity in development and progression of COPD.

In this study, COPD model was prepared by cigarette smoke exposure method. The experimental results showed that the animal model of COPD rats was made successfully from the behavioral activity and lung pathological changes. In lung pathological changes, the effect of low concentration of hydrogen is poor, the mechanism may be the antioxidant effect of hydrogen, low concentration of hydrogen can not resist the activation of the body to produce too much oxygen free radicals, and inhaled high concentration of hydrogen just neutralize this reaction. The mechanism may be the anti-inflammatory effect of hydrogen, which can reduce the release of inflammatory mediators and decrease the inflammatory factors to improve lung pathological status in COPD rats.

The experiment found that SIgA and PIgR decreased in COPD group, which may be due to the continuous stimulation of harmful gases, resulting in the destruction of bronchial mucosal structure, and decreased in the number of plasma cell, decreased IgA secretion(23) and reduced carrier PIgR. On the one hand, the composition and SC of synthetic carriers were reduced, on the other hand, the carrier was reduced, and finally the SIgA transported to the bronchial mucosal surface was reduced. Inhalation of high-concentration hydrogen significantly increased SIgA expression, and the effect was better than that of low-concentration hydrogen. This may be that the medium-high concentration of hydrogen stimulates plasma cell production, resulting in increased IgA secretion and PIgR delivery, so the SC of raw materials for synthetic SIgA will also increase, thus ameliorating the lack of airway mucosal surface.

The experimental study found that the expression of IL-4 was significantly decreased in COPD group, which may be caused by smoke stimulation Th cell dominant differentiation(24, 25). The expression of IL-4 and SIgAwas increased in Hh and Hm group, and the two closely related, so IL-4 may be one of the important factors affecting the expression of SIgA on the airway mucosal surface of rats with enhanced hydrogen. The hydrogen treatment group increased IL-5 expression to varying degrees. We hypothesized that hydrogen might increase IL-5 production by affecting Th2 cells, IL-5 promoting mucosal IgA production(26). A deficiency of the experiment is the lack of detection of Th2 cells, so whether the above views are true or not needs to be further improved to confirm.

We speculate that the other possible mechanism SIgA enhancement is that hydrogen can activate peripheral B cells. IL-40 participates in humoral immune response, and affects the number of IgA secretory cells, resulting in an increase in IgA production(16). IL-40, however, as a newly discovered factor in the last two years, few studies have been conducted to investigate whether IL-40 greatly affects SIgA expression as well as lung pathology. TGF-β family plays an important role in the development of human tissues and homeostasis. It regulates multiple cellular processes and is closely related to tissue remodeling in pulmonary fibrosis and emphysema(27). A number of studies have shown that TGF-β1 antagonists can ameliorate lung function, reduce collagen deposition, ameliorate lung pathology and airway remodeling(28, 29).

The results of this experiment show that hydrogen can effectively reduce the expression of TGF-β1, and statistics show that the expression of TGF-β1 is negatively correlated with the expression of SIgA. We believe that hydrogen interferes with lung pathology by down-regulating the expression level of SIgA.

## Conclusions

The model of COPD rats was successfully established after 4 months of smoke exposure. Through pathology observation and index measurement, hydrogen inhalation can ameliorate lung pathology injury in COPD rats. Hydrogen inhalation can significantly improve the expression of SIgA on the mucosal surface of COPD rats, which may be one of the mechanisms which hydrogen works on COPD pathogenesis. Hydrogen inhalation can improve the expression of IL-4, IL-5, and IL-40 factors in lung tissues, and reduce the expression of TGF-β1, these factors may play an important role in hydrogen changing the expression of SIgA on the mucosal surface of COPD rats.

## Acknowledgments

The authors would like to thank the Laboratory of Forensic Medicine of Hebei Medical University for its support.

## Author contributions

Wuzhuang Sun and Jiechao Wang planned and performed experiments, Yiping Ma and Yalei Zhao analysed data, and wrote the manuscript, Zhu Li and Mo Sun planned and performed experiments and analysed data.

## References

1. Mirza S, Clay RD, Koslow MA, Scanlon PD. COPD Guidelines: A Review of the 2018 GOLD Report. Mayo Clinic proceedings. 2018;93(10):1488–502.

2. Vestbo J, Hurd SS, Agusti AG, Jones PW, Vogelmeier C, Anzueto A, et al. Global strategy for the diagnosis, management, and prevention of chronic obstructive pulmonary disease: GOLD executive summary. American journal of respiratory and critical care medicine. 2013;187(4):347–65.

3. Xie S, Wang K, Zhang W, Xiao K, Yan P, Li Y, et al. Immunodeficiency in Patients with Acute Exacerbation of Chronic Obstructive Pulmonary Disease. Inflammation. 2018;41(5):1582–9.

4. Polosukhin VV, Richmond BW, Du RH, Cates JM, Wu P, Nian H, et al. Secretory IgA Deficiency in Individual Small Airways Is Associated with Persistent Inflammation and Remodeling. American journal of respiratory and critical care medicine. 2017;195(8):1010–21.

5. Ratajczak C, Guisset A, Detry B, Sibille Y, Pilette C. Dual effect of neutrophils on pIgR/secretory component in human bronchial epithelial cells: role of TGF-beta. Journal of biomedicine & biotechnology. 2010;2010.

6. Curtis JL. A Hairline Crack in the Levee: Focal Secretory IgA Deficiency as a First Step toward Emphysema. American journal of respiratory and critical care medicine. 2017;195(8):970–3.

7. Ladjemi MZ, Gras D, Dupasquier S, Detry B, Lecocq M, Garulli C, et al. Bronchial Epithelial IgA Secretion Is Impaired in Asthma. Role of IL-4/IL-13. American journal of respiratory and critical care medicine. 2018;197(11):1396–409.

8. Corthesy B. Role of secretory immunoglobulin A and secretory component in the protection of mucosal surfaces. Future microbiology. 2010;5(5):817–29.

9. Jaffar Z, Ferrini ME, Herritt LA, Roberts K. Cutting edge: lung mucosal Th17-mediated responses induce polymeric Ig receptor expression by the airway epithelium and elevate secretory IgA levels. Journal of immunology (Baltimore, Md: 1950). 2009;182(8):4507–11.

10. Liu DY, Jiang T, Wang S, Cao X. Effect of hyperoxia on pulmonary SIgA and its components, IgA and SC. Journal of clinical immunology. 2013;33(5):1009–17.

11. Bengoechea JA. Secretory IgA and COPD: a new kid on the block? American journal of respiratory and critical care medicine. 2011;184(3):285–7.

12. Du RH, Richmond BW, Blackwell TS, Jr., Cates JM, Massion PP, Ware LB, et al. Secretory IgA from submucosal glands does not compensate for its airway surface deficiency in chronic obstructive pulmonary disease. Virchows Archiv: an international journal of pathology. 2015;467(6):657–65.

13. Polosukhin VV, Cates JM, Lawson WE, Zaynagetdinov R, Milstone AP, Massion PP, et al. Bronchial secretory immunoglobulin a deficiency correlates with airway inflammation and progression of chronic obstructive pulmonary disease. American journal of respiratory and critical care medicine. 2011;184(3):317–27.

14. Richmond BW, Du RH, Han W, Benjamin JT, van der Meer R, Gleaves L, et al. Bacterial-derived Neutrophilic Inflammation Drives Lung Remodeling in a Mouse Model of Chronic Obstructive Pulmonary Disease. American journal of respiratory cell and molecular biology. 2018;58(6):736–44.

15. Catalan-Dibene J, McIntyre LL, Zlotnik A. Interleukin 30 to Interleukin 40. Journal of interferon & cytokine research: the official journal of the International Society for Interferon and Cytokine Research. 2018;38(10):423–39.

16. Catalan-Dibene J, Vazquez MI, Luu VP, Nuccio SP. Identification of IL-40, a Novel B Cell-Associated Cytokine. 2017;199(9):3326–35.

17. Wang C, Li J, Liu Q, Yang R, Zhang JH, Cao YP, et al. Hydrogen-rich saline reduces oxidative stress and inflammation by inhibit of JNK and NF-kappaB activation in a rat model of amyloid-beta-induced Alzheimer’s disease. Neuroscience letters. 2011;491(2):127–32.

18. Ohsawa I, Ishikawa M, Takahashi K, Watanabe M, Nishimaki K, Yamagata K, et al. Hydrogen acts as a therapeutic antioxidant by selectively reducing cytotoxic oxygen radicals. Nature medicine. 2007;13(6):688–94.

19. Manaenko A, Lekic T, Ma Q, Zhang JH, Tang J. Hydrogen inhalation ameliorated mast cell-mediated brain injury after intracerebral hemorrhage in mice. Critical care medicine. 2013;41(5):1266–75.

20. Iida A, Nosaka N, Yumoto T, Knaup E, Naito H, Nishiyama C, et al. The Clinical Application of Hydrogen as a Medical Treatment. Acta medica Okayama. 2016;70(5):331–7.

21. Liu X, Ma C, Wang X, Wang W, Li Z, Wang X, et al. Hydrogen coadministration slows the development of COPD-like lung disease in a cigarette smoke-induced rat model. International journal of chronic obstructive pulmonary disease. 2017;12:1309–24.

22. Wang L, Zhang B, Li Z, Li J, Liu Q, Sun W. Budesonide mitigates pathological changes in animal model Of COPD through reducing neutrophil elastase expression. International journal of clinical and experimental medicine. 2015;8(4):5227–35.

23. van Ginkel FW, Wahl SM, Kearney JF, Kweon MN, Fujihashi K, Burrows PD, et al. Partial IgA-deficiency with increased Th2-type cytokines in TGF-beta 1 knockout mice. Journal of immunology (Baltimore, Md: 1950). 1999;163(4):1951–7.

24. Chi Y, Di Q, Han G, Li M, Sun B. Mir-29b mediates the regulation of Nrf2 on airway epithelial remodeling and Th1/Th2 differentiation in COPD rats. Saudi journal of biological sciences. 2019;26(8):1915–21.

25. Sun J, Liu T, Yan Y, Huo K, Zhang W, Liu H, et al. The role of Th1/Th2 cytokines played in regulation of specific CD4 (+) Th1 cell conversion and activation during inflammatory reaction of chronic obstructive pulmonary disease. Scand J Immunol. 2018;88(1):e12674.

26. Sonoda E, Hitoshi Y, Yamaguchi N, Ishii T, Tominaga A, Araki S, et al. Differential regulation of IgA production by TGF-beta and IL-5: TGF-beta induces surface IgA-positive cells bearing IL-5 receptor, whereas IL-5 promotes their survival and maturation into IgA-secreting cells. Cellular immunology. 1992;140(1):158–72.

27. Saito A, Horie M, Nagase T. TGF-β Signaling in Lung Health and Disease. International journal of molecular sciences. 2018;19(8).

28. Kubiczkova L, Sedlarikova L, Hajek R, Sevcikova S. TGF-β - an excellent servant but a bad master. Journal of translational medicine. 2012;10:183.

29. Godinas L, Corhay JL, Henket M, Guiot J, Louis R, Moermans C. Increased production of TGF-β1 from sputum cells of COPD: Relationship with airway obstruction. Cytokine. 2017;99:1–8.

